# Construction and Analysis of Protein-Protein Interaction Network of Non-Alcoholic Fatty Liver Disease

**DOI:** 10.1101/2020.12.01.406215

**Authors:** Athina I. Amanatidou, George V. Dedoussis

## Abstract

Non-alcoholic fatty liver disease (NAFLD) is a disease with multidimensional complexities. Many attempts have been made over the years to treat this disease but its incidence is rising. For this reason, the need to identify and study new candidate proteins that may be associated with NAFLD is of utmost importance. Systems-based approaches such as the analysis of protein-protein interaction (PPI) network could lead to the discovery of new proteins associated with a disease that can then be translated into clinical practice. The aim of this study is to analyze the interaction network of human proteins associated with NAFLD as well as their experimentally verified interactors and to identify novel associations with other human proteins that may be involved in this disease. Computational analysis made it feasible to detect 77 candidate proteins associated with NAFLD, having high network scores. Furthemore, clustering analysis was performed to identify densely connected regions with biological significance in this network. Additionally, gene expression analysis was conducted to validate part of the findings of this research work. We believe that our research will be helpful in extending experimental efforts to address the pathogenesis and progression of NAFLD.

## 1. Introduction

The liver is a vital digestive organ which performs many essential body’s metabolic functions involving metabolism of lipids, bile acids, glucose and cholesterol [1]. Metabolic pathways do not operate independently within the liver; one pathway can heavily affect other pathways. The dysfunctional crosstalk of the hepatic pathways is a widespread health problem, responsible for about 2 million deaths worldwide each year [2]. The most common chronic liver disease worldwide is known as non-alcoholic fatty liver disease (NAFLD). It is an umbrella term which encompasses a spectrum of pathological conditions ranging from simple hepatic steatosis (SS) or non-alcoholic fatty liver (NAFL) to a more severe form nonalcoholic steatohepatitis (NASH), and NASH cirrhosis [3]. Although in the last decade, research advances demonstrate that NAFLD is a multisystem disease in which many complex processes are involved in its manifestation and development. In addition, growing number of studies demonstrates that NAFLD affects a variety of extrahepatic organs and regulatory pathways [4].

With the passage of time, NAFLD’s health and socio-economic influence is rising, and the annual health costs in the United States are greater than $103 billion [5]. Henceforth, its timely and precise diagnosis is very significant, considering that its prevalence has rapidly reached global epidemic proportions in both adults and children [6]. Most patients are asymptomatic and the diagnosis of the disease is random in most cases [7].

The medical community has centered on the causes of the disease over the past few decades, and the identification of new diagnostic markers (biomarkers). Nonetheless, the gold standard for NAFLD diagnosis remains the liver biopsy but this procedure is inefficient as a diagnostic tool due to its invasive, expensive and sometimes serious complications [8]. In the foreseeable future, the key to NAFLD diagnosis and treatment could be the “molecular signature” of each NAFLD patient [9].

The data that derived from omics technologies which feed precision medicine have a major contribution to this effort. An increasing number of technical advancements have, to date, produced a collection of many unused data as a whole. Therefore, it is necessary to move from single omics to multi-omics analysis, providing a broader window of its pathophysiology that scans different perspectives [9]. Network-based approaches integrate omics data such as protein-protein interaction (PPI) networks which are gaining ground in the scientific community as they provide valuable, quick and inexpensive tools for clarifying disease mechanisms and detecting new candidate disease-related proteins (or genes) [10].

Disease is rarely the result of an abnormality in a single gene but represents disruptions in the complex interaction network. Key biological factors that control the pathobiology of the disease are almost always the result of several pathobiological pathways interacting through an interconnected network [11]. Conventional methods which evaluate one gene or factor at a time have become less effective in tackling NAFLD’s multidimensional complexities [1]. Given the fact that NAFLD research mostly includes studies on human clinical and animal model trials [9], the analysis of PPI network could be an ally to uncover candidate biomarkers and pathological pathways, as well as potential therapeutic targets, contributing to the development of noninvasive diagnosis.

In the present study, a PPI network analysis was conducted to identify new candidate proteins that may be involved in NAFLD through performing topological analyses. Besides, clustering analysis of the PPI network was achieved to identify densely connected regions. In order to reveal insights into the molecular mechanisms of the network’s proteins, an enrichment analysis was performed. Moreover, an analysis of gene expression microarray data set was achieved to detect differential expressed genes (DEGs) between NAFLD samples and controls, as well as a pathway analysis of DEGs.

## 2. Methods

The research methodology used in this study includes the stages stated below. **Fig. 1** outlines the basic steps involved in the methodology.

**Fig. 1:**
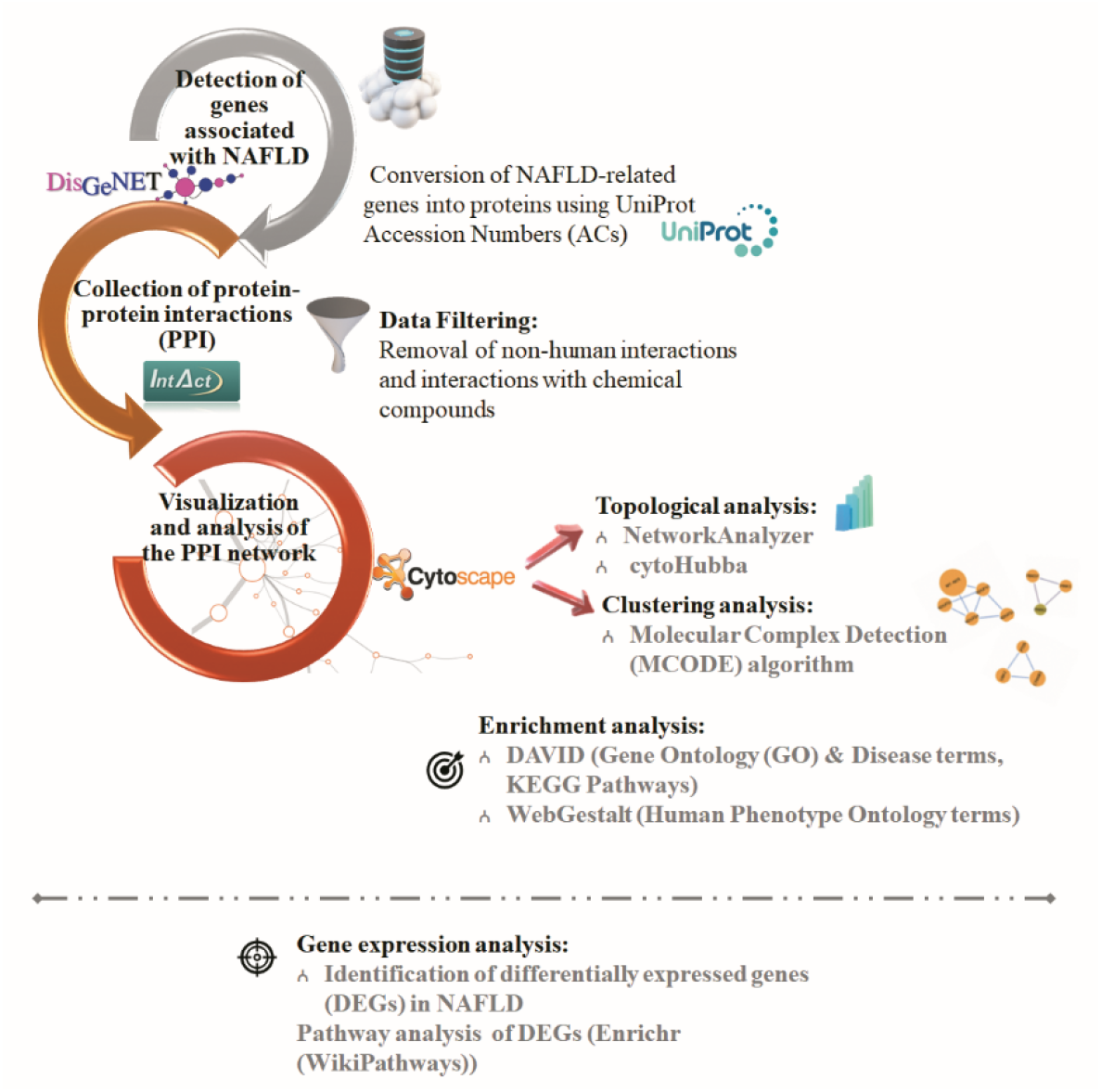
The schematic diagram of the research methodology.

### 2.1 Detection of genes associated with NAFLD

NAFLD and its subtype NASH have been queried using “Non-alcoholic Fatty Liver Disease” and “NASH - Nonalcoholic steatohepatitis” terms in a DisGeNET search panel which is a discovery platform containing one of the largest collections of genes and variants associated with human diseases [12]. All the NAFLD-related genes are either genetic associations or under/over expressed in the gene transcription levels or are present at low/high protein levels in patient’s plasma/serum. Eventually, the disease-related genes were manually confirmed for their association with NAFLD.

### 2.2 Collection of protein-protein interactions (PPI)

The NAFLD-related genes were then converted to proteins using UniProt Accession Numbers (ACs) via UniProt database [13]. A query was then conducted in IntAct [14], a molecular interaction database with highly curated data, using the ACs of the proteins, to retrieve all experimentally confirmed interactions of these proteins and their first neighbors. Interaction data were obtained in a MI-TAB 2.7 format file [15] in which any non-human interactions and interactions with chemical compounds were removed.

### 2.3 Visualization and analysis of the PPI network

Cytoscape (version) 3.7.2 software, a popular open source bioinformatics platform for the data integration and network analysis [16], was used to visualize and analyze the PPI network. In this network, every node corresponds to a protein and the edges represent interactions, where the latter were treated as undirected for this analysis. Additionally, browser-based web application was generated to visualize interactive networks via the CyNetShare tool (http://idekerlab.github.io/cy-net-share/). Links are provided in the legends of the respective figures.

Afterwards a **topological analysis** was conducted using the NetworkAnalyzer [17], a handy Cytoscape plugin, to estimate simple and complex topology parameters. The three important metrics – degree, betweenness and closeness centrality – were utilized to evaluate the importance of nodes in a network [10, 18]. *Hub* proteins were identified by their very high degree of connectivity. Proteins with high betweenness centrality, namely *bottlenecks,* are key connectors in the PPI network, controlling the flow of information within a network [19]. For the identification of proteins - from which the flow of information passes faster to other network’s proteins - are those with high closeness centrality, hereby referred to as *PHC (proteins with high closeness centrality)*[10]. The top scoring proteins corresponding to about the 5% of the network’s proteins were then selected for each of the three aforementioned network centralities. A Venn diagram was subsequently applied to identify **candidate NAFLD-related proteins** that were on the three high scoring protein lists but did not belong to the list of the NAFLD-related proteins.

Given the heterogeneous nature behind biological networks, it is advisable to use more than one approach to capture essential proteins. Therefore, a newly proposed method Maximal Clique Centrality (MCC) was estimated using the *cytoHybba* software [20], that has been proven for its great performance in predicting important proteins from the PPI network. The 10 top ranked proteins based on MCC algorithm were also identified as **candidate NAFLD-related proteins.**

Subsequently, Molecular Complex Detection (MCODE) algorithm was utilized to perform a **clustering analysis [21]**. The selection parameters were set as follows: MCODE scores>5, degree cut-off=2, node-score cut-off=0.2 and k-core=2.

Afterwards, an **enrichment analysis** was performed with the use of two bioinformatics tools, DAVID [22] and WebGestalt [23]. DAVID was used for functional enrichment analysis, disease association as well as pathway analysis and WebGestalt was utilized for human phenotype ontology (HPO) analysis. **Functional enrichment analysis** was applied to detect statistically significant overrepresented Gene Ontology (GO) [24] terms in the network. **Disease association analysis** was used to uncover the association of network’s proteins with disease terms from Gene Association Database (GAD) [25]. **Pathway analysis** was applied to detect the KEGG pathways from KEGG PATHWAY Database [26] and **HPO analysis**[27] used to detect the phenotype of network proteins’. P-value<0.05 was defined as statistical significance.

### 2.4 Gene expression data and pathway analyses of candidate NAFLD-related proteins

To detect differentially expressed genes (DEGs) in NAFLD compared to normal condition, the human gene expression data set GSE151158 [28] was downloaded from the Gene Expression Omnibus (GEO) (https://www.ncbi.nlm.nih.gov/geo/)[29], including 21 control liver samples, 40 NAFLD samples – 23 of which have NAFLD Activity Score (NAS) ≤ 3 and 17 have NAS ≥ 5. The analysis was performed through GEO2R [30] tool which applies *limma* (Linear Models for Microarray Analysis) [31] and *GEOquery*[32] R packages from the Bioconductor project. The data were log-transformed, and P-values were adjusted based on the Benjamini & Hochberg (False discovery rate, FDR) method for multiple testing. The significantly DEGs were defined with an adjusted P-value<0.05 and were then subjected to discover whether it contains any of the candidate NAFLD-related proteins resulting from the topological network analysis. The list of significantly DEGs were further analyzed against the WikiPathways [33] database by using the Enrichr[34] tool. P-value cutoff of 0.05 was selected to identify significantly enriched terms as well.

## 3. Results

### 3.1 Construction and analysis of NAFLD Interactome

The data set of NAFLD-related proteins is comprised of 254 proteins (**Supplementary Table 1**). They were then inserted into IntAct to collect their PPI, 226 of which have stored PPI data (**Supplementary Table 2**). Subsequently, the collected PPI data (**Supplementary Table 3**) were imported into Cytoscape 3.7.2 to construct a PPI network, refer to as ‘**NAFLD Interactome’**, comprising of 2624 proteins (nodes) and 20259 interactions (edges) (**Fig. 2**).

**Fig. 2:**
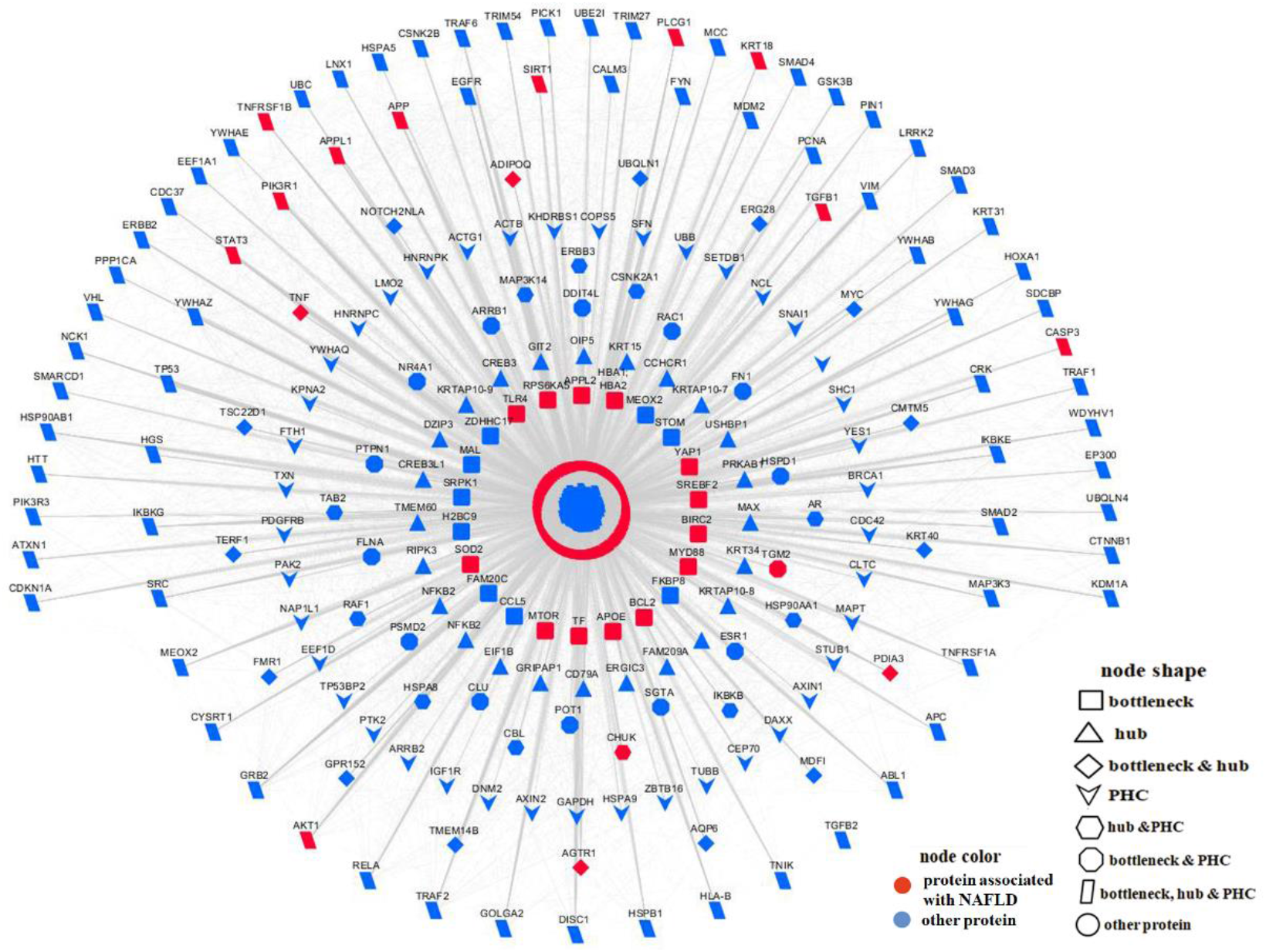
The NAFLD Interactome. A web visualization of this network is available at <u>/NAFLDInteractome</u>.

After conducting a **topological analysis** with the utilization of NetworkAnalyzer in NAFLD Interactome, important information regarding the network’s topology and the biological value of its proteins was revealed. The network’s density (show how sparse/dense is a network) is estimated as 0.006, a value lower than 0.1, which denotes that the NAFLD Interactome is a sparsely connected network, as other biological networks [35]. The clustering coefficient, the propensity of the network to grouped into clusters, is measured as 0.110 and the characteristic path length (CPL) [36] is 3.285.

The node degree distribution *P(k)*[37], follows the power-law P(k) =*Ak^−γ^*, where A is constant and γ is the degree exponent. In our case, the distribution is of the following form:

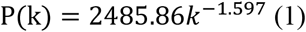

PPI networks are scale-free and its main feature is that they follow the power law node degree distribution [38]. Since this network also follows the power law distribution; it is characterized by a small number of highly connected proteins, while the majority of the other proteins have few interactions with others [37].

To quantify the importance of network’s proteins, metrics for the degree, betweenness and closeness centrality were applied for all NAFLD interactome’s proteins. Specifically, the proteins were ranked based on the three afore mentioned centrality measures and then the top 5% of the network’s proteins with the highest values were chosen. Considering the overlapping proteins among the protein lists of each network centrality, a total of 208 proteins were finally selected (**Supplementary Table 4**). Particularly, in the NAFLD Interactome, 25 proteins are *hubs* (**Fig. 2**, triangles), 22 proteins are *bottlenecks* (**Fig. 2**, rectangles), 17 proteins are *hubs* and *bottlenecks* (**Fig. 2**, diamonds), 40 proteins are *PHCs* (**Fig. 2**, V-shaped nodes), 11 proteins are *hubs* and *PHCs* (**Fig. 2**, hexagons), 14 proteins are *bottlenecks* and *PHCs* (**Fig. 2**, octagons), and 79 proteins are *hubs*, *bottlenecks* and *PHCs* (**Fig. 2**, parallelograms). It is noteworthy that 30 NAFLD-related proteins play an essential role in the NAFLD Interactome.

The **enrichment analysis** in NAFLD Interactome (2624 proteins) was performed to uncover the role of the network’s proteins (more details are given in **Supplementary Tables 5-8**). Among of the most statistically significant over-represented **GO terms** are the following: negative (GO:0043066) (P-value: 5.22E-37) and positive regulation of apoptotic process (GO:0043065) (P-value: 3.78E-34), positive regulation of transcription from RNA polymerase II promoter (GO:0045944) (P-value: 4.19E-34) and inflammatory response (GO:0006954) (P-value: 9.55E-28).

The **KEGG pathways terms** in which most proteins were found to be involved are pathways in cancer (hsa05200) (P-value: 4.11E-41), PI3K-Akt signaling pathway (hsa04151) (P-value: 1.15E-25), proteoglycans in cancer (hsa05205) (P-value: 1.11E-29), MAPK signaling pathway (hsa04010) (P-value: 5.67E-18) and focal adhesion (hsa04510) (P-value: 3.89E-24). The **disease association analysis** shows that type 2 diabetes (P-value: 1.91E-52), chronic kidney failure (P-value: 3.90E-38), Alzheimer’s disease (P-value: 9.41E-23), lung (P-value: 5.84E-53), bladder (P-value: 1.12E-48) and breast (P-value: 9.83E-54) cancer, as well as multiple sclerosis (P-value: 8.62E-27) and schizophrenia (P-value: 4.56E-17) are among of the numerous identified disease terms. Moreover, several phenotypic abnormalities were identified from **HPO analysis** including abnormality of the digestive system (HP: 0025031) (P-value: 5.23E-08), metabolism/homeostasis (HP: 0001939) (P-value: 2.26E-07), cardiovascular system (HP: 0001626) (P-value: 2.53E-04), skin morphology (HP: 0011121) (P-value: 1.96E-07) and immune system (HP: 0002715) (P-value: 6.89E-10).

Two different approaches were applied to identify candidate NAFLD-related proteins, as previously described in the Methods section. In the first approach, in order to find which proteins are present in the list of 79 high scoring proteins (hubs, bottlenecks and PHCs) and already associated with NAFLD, the list of high scoring proteins was combined with the list of 226 NAFLD-related proteins using Venn diagram. Thusly, 68 proteins were recognized as belonging only to the list of high scoring proteins, called candidate NAFLD-related proteins (**Table 1a)**. In the second approach, the 10 top-ranked proteins were found applying MCC algorithm, which are given in **Table 1b**. While CLOCK belongs to the list of 226 NAFLD-related proteins, the remaining 9 proteins were identified as candidate NAFLD-related proteins.

**Table 1a:**
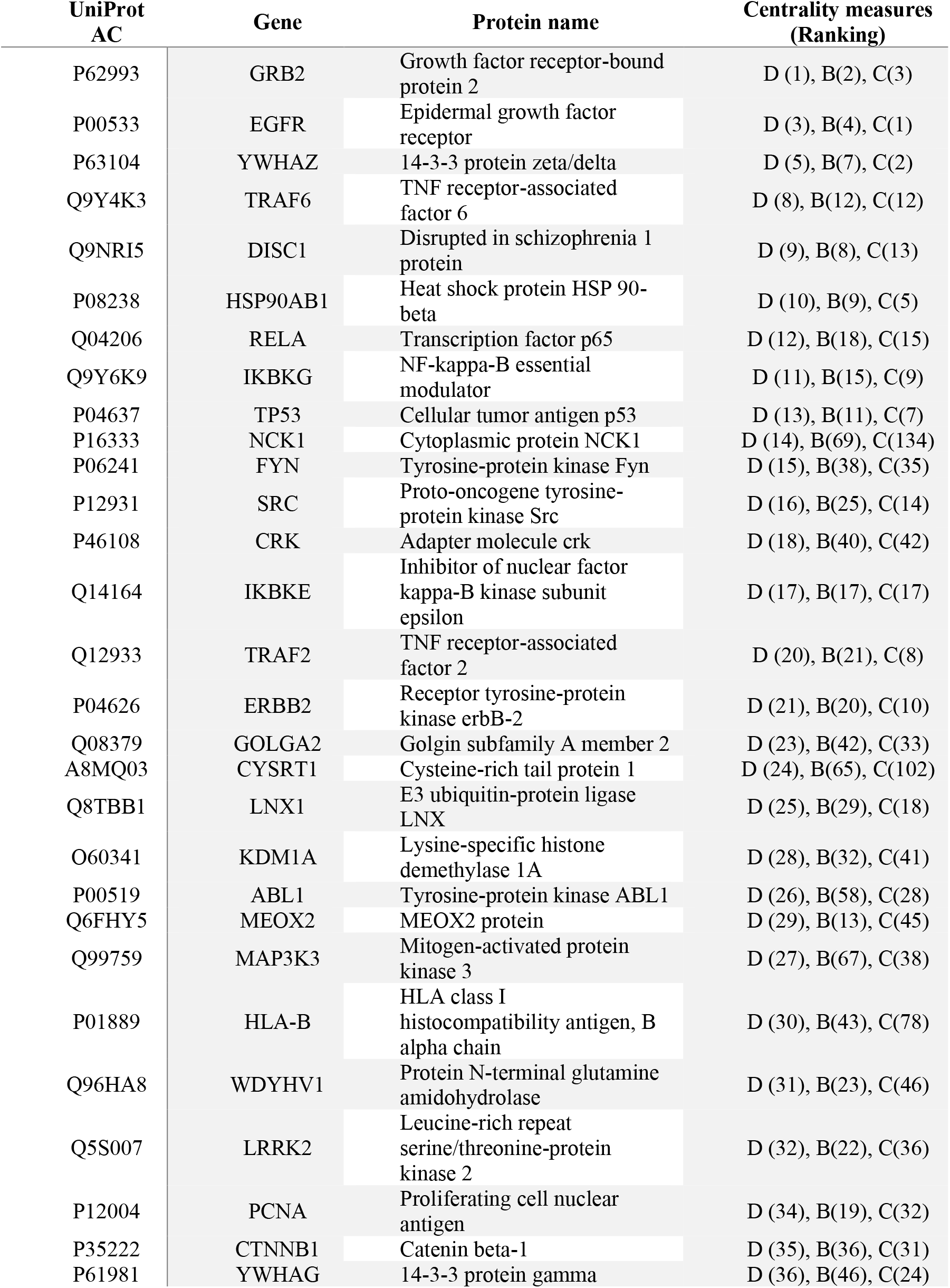
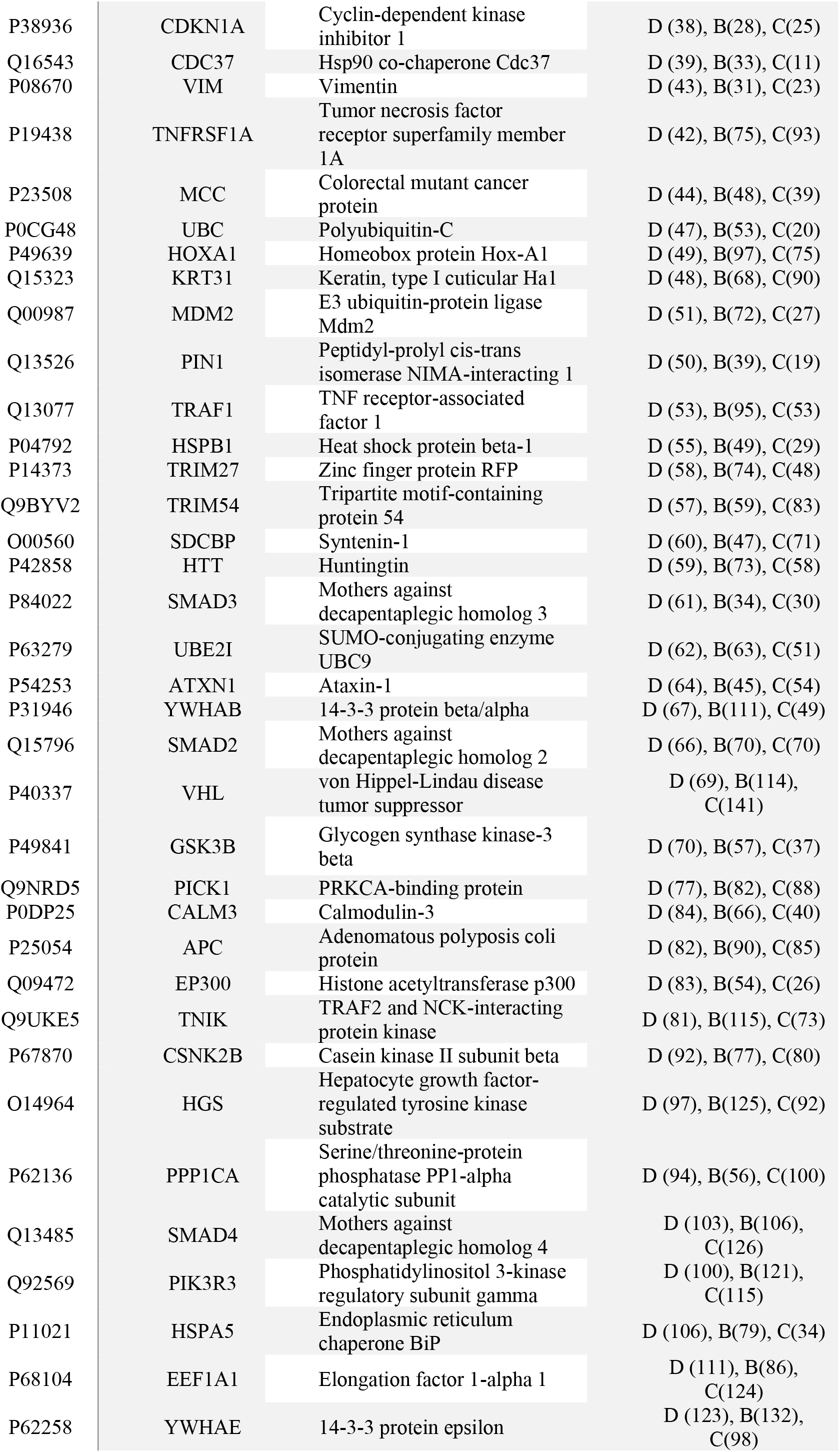
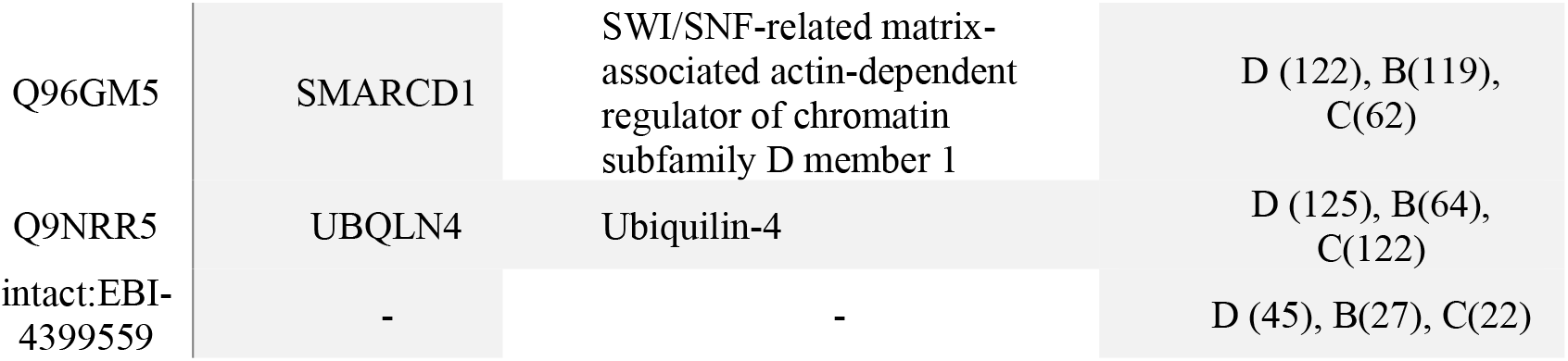
Identification of candidate NAFLD-related proteins. The column “Centrality measures” shows the proteins’ ranking in Degree-D, Betweenness-B and Closeness-C network centrality measures. The rank of each protein is given inside the parenthesis of the corresponding centrality measure in the top 140 rankings (approximately the top 5% of the network’s proteins).

**Table 1b:** Identification of candidate NAFLD-related proteins. The 10 top-ranked proteins based on MCC method in NAFLD Interactome. CLOCK protein, highlighted in bold, is already in the list of NAFLD-related proteins.

| UniProt AC | Gene | Protein name |
| --- | --- | --- |
| **O15516** | **CLOCK** | **Circadian locomoter output cycles protein kaput** |
| Q9UKL0 | RCOR1 | REST corepressor 1 |
| Q9NNX1 | TUFT1 | Tuftelin |
| Q96BD5 | PHF21A | PHD finger protein 21A |
| O43482 | OIP5 | Opa-interacting protein 5 |
| Q86Y13 | DZIP3 | E3 ubiquitin-protein ligase DZIP3 |
| Q9NP66 | HMG20A | High mobility group protein 20A |
| O95619 | YEATS4 | YEATS domain-containing protein 4 |
| Q96JG6 | VPS50 | Syndetin |
| Q567U6 | CCDC93 | Coiled-coil domain-containing protein 93 |

The results of the **enrichment analysis** of candidate NAFLD-related proteins are shown in **Supplementary Table 9**.

### 3.2 Clustering and enrichment analysis

#### Clustering analysis

The base of this study is the NAFLD Interactome, a large interconnected network with interactive embedded subnetworks. Hence, with a valuable applying of clustering analysis via MCODE algorithm, the detection of 6 clusters with MCODE score>5 was achieved (**Fig. 3**). The first cluster (MCODE score=29.655) consists of 30 proteins, including 1 NAFLD-related protein: CLOCK (**Fig. 3, 1^st^ Cluster-red node**). It is of utmost importance for our analysis to note that 9 of which are candidate NAFLD-related proteins: **RCOR1, TUFT1, PHF21A, OIP5, DZIP3, HMG20A, YEATS4, VPS50 and CCDC93**(**Fig. 3, 1^st^ Cluster-magenta nodes**). Also, the second cluster (MCODE score=15.412) integrates 18 proteins 2 of which are candidate NAFLD-related proteins: **HOXA1 and CYSRT1**(**Fig. 3, 2^nd^ Cluster-magenta nodes**).

**Fig. 3:**
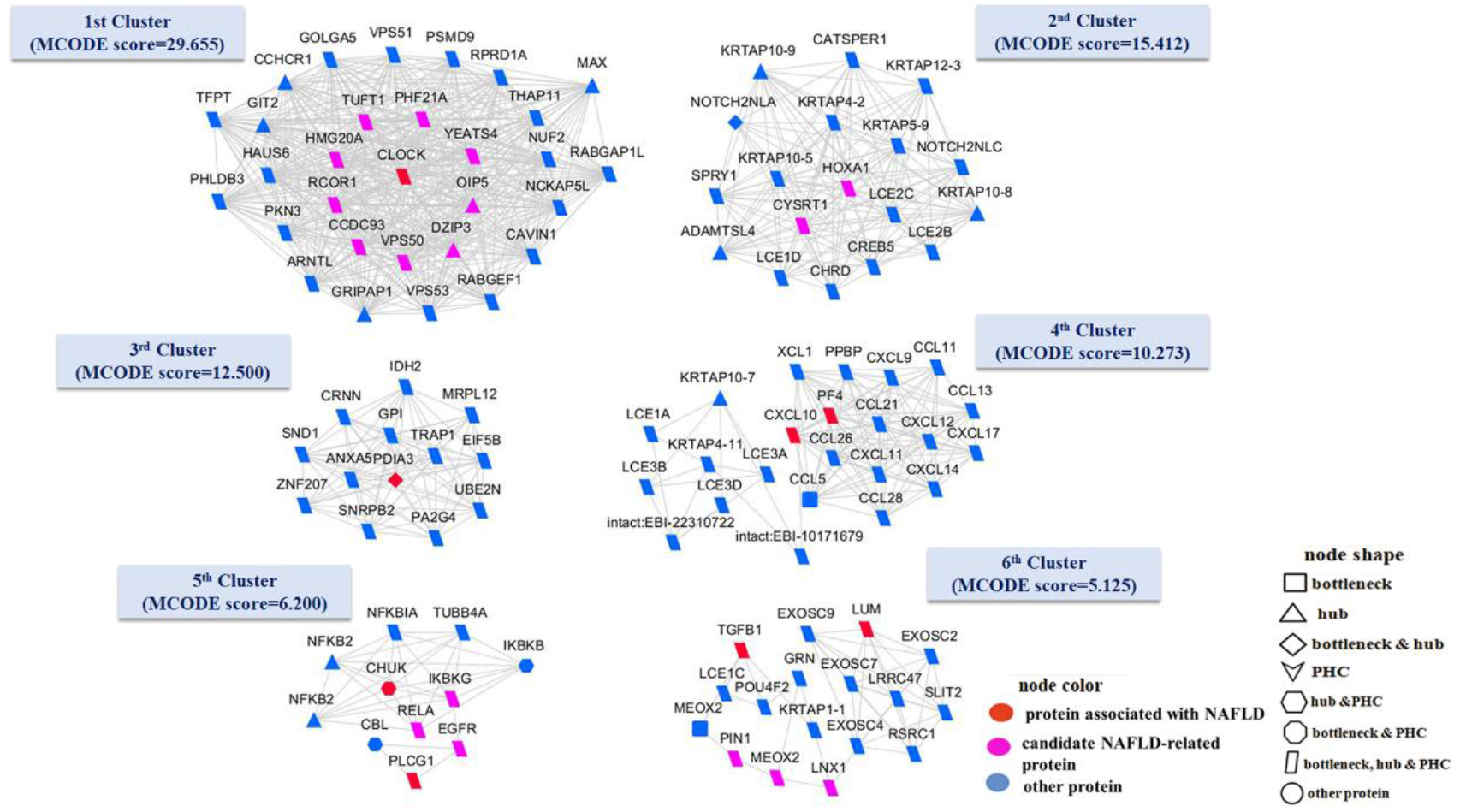
Clustering analysis of the NAFLD Interactome. A web visualization of this network is available at/<u>ClusteringAnalysisNAFLDInteractome</u>.

Subsequently, the third (MCODE score=12.500) and fourth (MCODE score=10.273) cluster comprise of 13 and 23 proteins, respectively, containing 1 NAFLD-related protein: PDIA3 (**Fig. 3, 3^rd^ Cluster-red node**) and 2 NAFLD-related proteins: CXCL10 and PF4 (**Fig. 3, 4^th^ Cluster-red nodes**), correspondingly. The fifth cluster (MCODE score=6.200) integrates 11 proteins, 2 of which are NAFLD-related proteins: CHUK and PLCG1 (**Fig. 3, 5^th^ Cluster-red nodes**) and 3 are candidate NAFLD-related proteins: **RELA, IKBKG and EGFR**(**Fig. 3, 5^th^ Cluster-magenta nodes**). Finally, the sixth cluster (MCODE score=5.125) encompasses 17 proteins, involving 2 NAFLD-related proteins: LUM and TGFB1 (**Fig. 3, 6^th^ Cluster-red nodes**) and 3 candidate NAFLD-related proteins: **MEOX2, LNX1 and PIN1**(**Fig. 3, 6^th^ Cluster-magenta nodes**).

#### Functional enrichment analysis

GO terms were detected for each cluster. Specifically, BP terms could be extracted for the 1st, 2nd, 4th, 5th and 6th clusters (**Supplementary Table 10**), while the MF and CC terms are identified for all clusters (**Supplementary Table 11-12**).

#### Pathway analysis

The pathway analysis brings to light information regarding the common pathways in which each cluster’s proteins partake. Results were detected for all clusters except for the 2^nd^ cluster. Circadian rhythm (hsa04710) (P-value: 0.0223) was found present in the 1^st^ cluster. Chemokine signaling pathway (hsa04062) (P-value: 2.72E-21) and cytokine-cytokine receptor interaction (hsa04060) (P-value: 9.73E-20) dominated in the 4^th^ cluster. Moreover, the majority of 5 ^th^ cluster’s proteins were found to be involved in epithelial cell signaling in Helicobacter pylori infection (hsa05120) (P-value: 5.58E-11) and NF-kappa B signaling pathway (hsa04064) (P-value: 2.80E-10). Finally, only RNA degradation (hsa03018) (P-value: 4.57E-05) was detected in 6^th^ cluster. No results were returned for the 2^nd^ cluster. More details of pathway analysis are given in **Supplementary Table 13**.

#### Disease association analysis

Statistically significant disease terms were retrieved for each cluster, although no results were detected for the 2^nd^ cluster (**Supplementary Table 14**). Interestingly, depression (P-value: 0.0193) and sleep disorders (P-value: 0.0368) are associated with the 1^st^ cluster’s proteins. Acquired immunodeficiency syndrome (P-value: 0.0147) is the only statistically significant term of the 3^rd^ cluster and respiratory syncytial virus bronchiolitis (P-value: 3.77E-11) is highly related to the 4^th^ cluster’s proteins. Also, rheumatoid arthritis (P-value: 1.86E-09) and benzene haematotoxicity (P-value: 3.82E-07) are among the highly statistical terms associated with proteins of the 5^th^ cluster. Lastly, vesico-ureteral reflux (P-value: 0.0055) was found to be the most statistically significant term of the 6^th^ cluster’s proteins.

#### HPO analysis

Phenotypic abnormality terms are detected for all clusters apart from 4^th^ cluster. Please refer to **Supplementary Table 15**for more details.

### 3.3 Gene expression data and pathway analyses of candidate NAFLD-related proteins

#### Identification of DEGs

A gene expression analysis was performed to detect DEGs that were differentially expressed between 23 NAFLD-NAS ≤ 3 samples and 21 controls (NAFLD-NAS ≤ 3 vs. Controls), between 17 NAFLD-NAS ≥ 5 samples and 21 controls (NAFLD-NAS ≥ 5 vs. Controls), and between 40 NAFLD samples and 21 controls (NAFLD-all vs. Controls). A total of 55 DEGs, 249 DEGs and 223 DEGs were identified between NAFLD-NAS ≤ 3 vs. Controls, NAFLD-NAS ≥ 5 vs. Controls and NAFLD-all vs. Controls, respectively. In accordance with our results, **TRAF1, HLA-B, IKBKE and SRC** are the genes that previously were identified as candidate NAFLD-related proteins and were also found as differentially expressed between NAFLD-NAS ≤ 3, NAFLD-NAS ≥ 5, NAFLD-all and Controls. Likewise, **TRAF2, CDKN1A and TP53** were found common between NAFLD-NAS ≥ 5, NAFLD-all and Controls. Please refer to the **Supplementary Table 17**for further details.

#### Pathway analysis of DEGs

In NAFLD-NAS ≤ 3 vs. Controls, NAFLD-NAS ≥ 5 vs. Controls and NAFLD-all vs. Controls contrast groups, DEGs were significantly enriched in 93, 186 and 185 pathways, respectively (**Supplementary Tables 18 A-C**). The top 10 enriched pathways of DEGs that were most statistically significant between NAFLD-NAS ≤ 3, NAFLD-NAS ≥ 5 and Controls are shown in **Table 2**. Interestingly, **IKBKE** is involved in several pathways such as regulation of toll-like receptor signaling pathway and RIG-I-like Receptor Signaling; **SRC** is implicated in Fibrin Complement Receptor 3 Signaling Pathway and Viral Acute Myocarditis; **HLA-B** is enriched in Allograft Rejection and Type II interferon signaling; **TRAF1, TRAF2 and TP53** are associated with apoptosis; **CDKN1A, SRC and TP53** are implicated in Senescence and Autophagy in Cancer.

**Table 2:** The top 10 most significantly enriched pathways of DEGs between NAFLD-NAS ≤ 3, NAFLD-NAS ≥ 5 and Controls. The genes that previously identified as candidate NAFLD-related proteins are highlighted in bold.

| Term | P-value | Count | Genes |
| --- | --- | --- | --- |
| **NAFLD-NAS $\leq 3$** |  |  |  |
| Regulation of toll-like receptor signaling pathway (WP1449) | 4.25E-12 | 10 | CXCL10, CXCL9, CASP8, SYK, IRF7, SPP1, LY96, CD14, TNF, **IKBKE** |
| Fibrin Complement Receptor 3 Signaling Pathway (WP4136) | 7.73E-12 | 7 | CXCL10, SYK, **SRC**, ITGB2, LY96, CD14, TNF |
| Toll-like Receptor Signaling Pathway (WP75) | 9.32E-12 | 9 | CXCL10, CXCL9, CASP8, IRF7, SPP1, LY96, CD14, TNF, **IKBKE** |
| Apoptosis (WP254) | 7.14E-11 | 8 | CASP8, CASP3, CASP1, IRF7, BAX, FAS, **TRAF1**, TNF |
| Viral Acute Myocarditis (WP4298) | 7.14E-11 | 8 | CASP8, **SRC**, CASP3, ITGB2, CASP1, BAX, NOD2, TNF |
| Allograft Rejection (WP2328) | 1.59E-07 | 6 | CXCL9, CASP8, CASP3, **HLA-B**, FAS, TNF |
| Nanomaterial induced apoptosis (WP2507) | 2.40E-07 | 4 | CASP8, CASP3, FAS, BAX |
| RIG-I-like Receptor Signaling (WP3865) | 6.36E-07 | 5 | CXCL10, CASP8, IRF7, TNF, **IKBKE** |
| Type II interferon signaling (IFNG) (WP619) | 3.16E-06 | 4 | CXCL10, CXCL9, **HLA-B**, PSMB9 |
| Amyotrophic lateral sclerosis (ALS) (WP2447) | 3.52E-06 | 4 | CASP3, CASP1, BAX, TNF |
| **NAFLD-NAS $\geq 5$** |  |  |  |
| Allograft Rejection (WP2328) | 1.05E-38 | 32 | CD86, CXCL9, ABCB1, CD80, PRF1, CXCL13, HLA-DMB, **HLA-B**... |
| Regulation of toll-like receptor signaling pathway (WP1449) | 4.69E-26 | 28 | CD86, CXCL9, CD80, LY96, TNFAIP3, TNF, CASP8, CCL5, CCL4, **IKBKE**... |
| Viral Acute Myocarditis (WP4298) | 6.47E-25 | 23 | TGFB1, **SRC**, STAT1, CD80, ITGB2, CXCR4, NOD2... |
| Toll-like Receptor Signaling Pathway (WP75) | 4.24E-24 | 24 | CD86, CXCL9, STAT1, CD80, LY96, TNF, **IKBKE**, TLR3... |
| Ebola Virus Pathway on Host (WP4217) | 4.95E-19 | 22 | **HLA-B**, ICAM3, HLA-C, HLA-A, NFKB2, HLA-DMA, HLA-DMB, IRF7, HLA-DPB1, **IKBKE**... |
| Chemokine signaling pathway (WP3929) | 1.02E-16 | 22 | CCR1, CX3CR1, CXCL9, CCL22, CCL20, STAT1... |
| Human Complement System (WP2806) | 3.49E-15 | 17 | SELPLG, C1R, ITGB2, PLAUR, C8A, C2, C5... |
| Apoptosis (WP254) | 5.80E-15 | 16 | **TRAF2**, **TRAF1**, TNF, CASP8, CASP10, **TP53**... |
| Senescence and Autophagy in Cancer (WP615) | 1.39E-14 | 17 | **CDKN1A**, TGFB1, **SRC**, ATG10, IFI16, IL1B, **TP53**... |
| T-Cell antigen Receptor (TCR) Signaling Pathway (WP69) | 1.82E-14 | 16 | MAP4K1, CD83, TGFB1, PRKCD, NFATC1... |

## 4. Discussion

PPI networks are widely accepted for their valuable contribution to the identification of candidate disease-related proteins in several diseases such as hepatocellular carcinoma, blood-cell targeting autoimmune diseases, breast cancer, etc [10, 39, 40]. In the present study, a topological analysis of the NAFLD Interactome was conducted by applying two different approaches (as presented throughout the Methods section), thusly a **total of 77 candidate NAFLD-related proteins were identified**. Surprisingly, about 50% of these proteins are previously verified in human and animal studies, as well as in other bioinformatics studies regarding their implication in NAFLD and in liver-related manifestations. The validation of our results through literature, which are described bellow, shows that the approach followed in this study is effective in identifying candidate NAFLD-related proteins. Therefore, the remaining unconfirmed proteins should be further investigated for their possible association with NAFLD.

The findings of our literature survey confirmed the implication of the following: **HSP90AB1** has been suggested as a possible biomarker in overweight and obese children with NAFLD [41]; **HLA-B**[42], **CTNNB1**[43] and **HSPA5**[44] are found to be abnormally expressed in NAFLD patients; **CDKN1A**polymorphism is associated with the development of human NAFLD [45]; **TRAF1** has been also detected in NAFLD patients [46]; **HSPB1**phosphorylation site has been differed between NAFLD cohorts [47]; **SMAD4** was overexpresed in NASH patients [48]; **SMAD2/3**phosphorylation and nuclear translocation documented in the liver of NASH patients[49]; **RELA** is well-known to cause inflammatory responses in NAFLD [50]; **PIK3R3** has been proposed as an effective candidate target for the development of NAFLD [51]; **GSK3B** inhibition has been proposed as a possible therapeutic target to manipulate the NAFLD [52].

Remarkably, our findings are in aggreement with previous animal studies as mentioned below: **EGFR** inhibition has been proved to attenuate NAFLD in obese mice model, playing an essential role in NAFLD as a possible therapeutic target [53]; **TP53** inhibition in a NAFLD mice model resulting in decreased steatosis and liver injury [54]; **PIN1** was essentially involved in NASH development in a rodent model [55]; **SMAD3** overexpression was identified in the liver of monkeys with simple steatosis (SS) and fibrosing NASH [56]; **KDM1A**elevated expression was found in NASH-related hepatocarcinogenesis in a mice model [57]; **EEF1A1** inhibition has been shown to reduce lipotoxicity in obese mice with NAFLD [58]; **TNFRSF1A** has been identified as a potentially effective target factor to prevent the attenuation of SS progression to a more complex phenotype with many NASH features in a mice model [59]; **IKBKE** has been found to specifically expressed in hepatic stellate cells (HSCs) in which inhibition by amlexanox in a NAFLD mice model resulted in improved insulin signal pathway in hepatocytes [60]; **FYN** is implicated in fatty acid oxidation and hepatic steatosis development under chronic ethanol intake in mice model [61]; the increased expression of **VIM** has been found during hepatic steatosis development to NASH in mice, suggesting it as a valuable prognostic factor of liver disease severity [62]; **VIM** and **MAP3K3** were identified upregulated by decreased liver miR-122, possibly contributing in NASH-induced hepatic fibrosis in mice [63]; **ABL1** is implicated in axis which regulates a murine hepatic steatosis, serving as candidate anti-steatosis target [64]; **EP300** inhibition could be effective in hepatic steatosis in mice [65].

In light of the literature review, our results seem to be promising regarding their possible implication in NAFLD development and progression. Recently, **YWHAZ** has been defined as a new regulator of several genes which are dysregulated in NAFLD development [66]. Remarkably, the genetic dysfunction of **MDM2** in adipocytes activates apoptotic and senescent **TP53**-mediated programs causing lipodystrophy and its related several metabolic diseases such as NAFLD [67]. Also, **VHL**disruption resulted in significant lipid accumulation, hepatic inflammation and fibrosis in the liver [68]. Lately, **SRC** has been found upregulated during the hepatic HSCs activation and liver fibrosis [69]. Also, **IKBKG**(or NEMO) deletion in liver parenchymal cells results in steatohepatitis and hepatocellular carcinoma [70]. Furthermore, **GRB2**suppression has been shown to improve hepatic steatosis, glucose metabolism, apoptosis and oxidative stress [71]. Moreover, the decreased expression of **SMARCD1** activates lipid accumulation and cellular senescence, denoting its preventative role regarding lifestyle-related diseases [72]. The phospho-**UBE2I** has been suggested to potentially enhance NF-kB signaling, revealing a possible new mechanism that deregulates inflammatory signaling of the liver [73]. The **GOLGA2** inhibition is found to induce fibrosis with autophagy in the liver and lung of mice [74]. **ERBB2**(also known as HER2) is closely linked to many enzymes, e.g. fatty acid synthase, which play essential regulatory roles in lipid metabolism or lipogenic pathways [75] and its hepatic expression has been identified in liver diseases [76, 77]. Remarkably, the hepatic gene expression of **SDCBP** has been found differentially expressed in steatotic liver [78]. Also, **CDC37** was defined with a modulatory role of INK4A activity in rat hepatic carcinogenesis and human hepatic cancer [79].

Interestingly, several studies applying bioinformatics analyses are in consistensy with our findings, revealing the possible implication of **UBQLN4**[80], **UBC**[81] and **PCNA**[82] in NAFLD development as potential biomarkers. Likewise, a bioinformatics analysis in a PPI network of steatosis highlights **CRK** and **MDM2** among of the top 10 important genes [83].

It is a well-known fact that disease-related proteins are clustered together and are also centrally located within a network [84]. As demonstrated from our results, the identified candidate NAFLD-related proteins: **RCOR1, TUFT1, PHF21A, OIP5, DZIP3, HMG20A, YEATS4, VPS50, CCDC93 (Fig. 3, 1st Cluster-magenta nodes), RELA, IKBKG, EGFR (Fig. 3, 5th Cluster-magenta nodes), MEOX2, LNX1 and PIN1 (Fig. 3, 6th Cluster-magenta nodes),** are found in the same clusters with already known NAFLD-related proteins, enhancing their potential implication in NAFLD. Notably, **RELA, IKBKG, EGFR** and **PIN1**, as already mentioned, are literally confirmed for their possible association with NAFLD.

Worthwhille to mention that the 7 candidate NAFLD-related proteins: **TRAF1, TRAF2, HLA-B, IKBKE, SRC, CDKN1A** and **TP53** are validated through the gene expression analysis. At first glance, this will probably not seem very prominent but it does show that the network approach followed in this study is complementary to gene expression analysis by identifying more candidates associated with NAFLD that would otherwise not be detected. After performing pathway analysis of DEGs, **IKBKE** was found to be involved in toll-like receptor signaling pathway that play an important role in the NAFLD development [85]. Moreover, **TRAF1, TRAF2** and **TP53** are implicated in apoptosis which seems to be important in NAFLD and NASH progression [86]. Reportedly, **CDKN1A, SRC** and **TP53** are participated in senescence and autophagy in cancer. Interestingly, considerable associations have been established between regulation of autophagy and obesity-related liver complications, NAFLD [87]. It is important to mention that human clinical studies revealed the association of senescence with NAFLD [88]. Thereby, the aforementioned genes might play pivotal roles in the development and progression of NAFLD via regulating the pathways involved in this disease.

The **enrichment analysis** of the NAFLD Interactome was performed to examine the functional and biological interactions among the proteins, as well as to uncover their associations with diseases and several phenotypic abnormalities in human. Pathway analysis revealed that proteins are significantly enriched among others in pathways in cancer, PI3K-Akt signaling pathway, proteoglycans in cancer, MAPK signaling pathway and focal adhesion. It has been demonstated that PI3K-Akt and MAPK signaling pathways have been shown to be involved in NAFLD [89, 90]. Moreover, focal adhestion kinase regulates the activation of HSCs and liver fibrosis [91]. Interestingly, in the wound healing response, focal adhesion and proteoglycans in cancer pathways are implicated. As stated by other research works, these wound healing and cell migration pathways have been shown to be dysregulated in NASH leading to fibrosis [92]. Disease association analysis showed that proteins are associated with a number of diseases such as type 2 diabetes [93], chronic kidney failure [94], Alzheimer’s disease [95], multiple sclerosis, schizophrenia [96], lung, bladder [97] and breast cancer [98], most of which are associated with NAFLD. Also, the phenotypic abnormalities of proteins such as those of digestive system, metabolism/homeostasis, cardiovascular system, skin morphology and immune system are linked with NAFLD [99–102].

In conclusion, applying a systemic approach to this study, we were able to identify **77 candidate NAFLD-related proteins,** out of which 41 *(HSP90AB1, HLA-B, CTNNB1, HSPA5, CDKN1A, SMAD4, SMAD2, SMAD3, TRAF1, HSPB1, RELA, PIK3R3, GSK3B, VHL, SRC, EGFR, TP53, PIN1, KDM1A, EEF1A1, UBQLN4, UBC, PCNA, CRK, MDM2, VIM, MAP3K3, TNFRSF1A, YWHAZ, IKBKG, FYN, ABL1, GRB2, SMARCD1, UBE2I, GOLGA2, IKBKE, EP300, ERBB2, SDCBP,CDC37)* are confirmed through literature searches. The novelty of our findings lies in the remaining 36 proteins *(TRAF6, DISC1, NCK1, TRAF2, CYSRT1, LNX1, MEOX2, WDYHV1, LRRK2, YWHAG, MCC, HOXA1, KRT31, TRIM27, TRIM54, HTT, ATXN1, YWHAB, PICK1, CALM3, APC, TNIK, CSNK2B, HGS, PPP1CA, YWHAE, RCOR1, TUFT1, PHF21A, OIP5, DZIP3, HMG20A, YEATS4, VPS50, CCDC93, intact:EBI-4399559)*that could may be involved in NAFLD. It should be pointed out that the implementation of clustering analysis revealed the importance of 15 candidate NAFLD-related proteins in NAFLD *(RCOR1, TUFT1, PHF21A, OIP5, DZIP3, HMG20A, YEATS4, VPS50, CCDC93, RELA, IKBKG, EGFR, MEOX2, LNX1 and PIN1)* in light of the fact that are clustered together with known NAFLD-related proteins. Also, 9 of which *(RCOR1, TUFT1, PHF21A, OIP5, DZIP3, HMG20A, YEATS4, VPS50 and CCDC93)*had not been published before in other research works. Noteworthy, we subsequently achieved via gene expression analysis the verification of 7 candidate NAFLD-related proteins: *TRAF1, TRAF2, HLA-B, IKBKE, SRC, CDKN1A and TP53,* while TRAF2 is one of the proteins that has not been found previously in the literature. Several of the results obtained in the present study are also reported by many other studies, as outlined in the Discussion section of this manuscript. We hope that our research will serve as a base for further experimental works.

## Acknowledgements

The authors thank the Harokopio University of Athens for use of premises and equipment.

The research work was financially supported by the Hellenic Foundation for Research and Innovation (HFRI) under the HFRI PhD Fellowship grant (Fellowship Number: 1529).

## CRediT author statement

**Athina I. Amanatidou:** Conceptualization, Methodology, Validation, Formal analysis, Data Curation, Investigation, Visualization, Writing - Original Draft. **George. V. Dedoussis:**Supervision, Writing - Review & Editing.

## Abbreviations

NAFLD: non-alcoholic fatty liver disease
NASH: nonalcoholic steatohepatitis
PMIDs: PubMed IDs
PPI: Protein-protein interaction
MCODE: Molecular Complex Detection
PHC: Proteins with high closeness centrality
HPO: Human Phenotype Ontology
DEGs: Differentially expressed genes
CPL: Characteristic path length
NAS: NAFLD Activity Score
HSP90AB1: Heat shock protein HSP 90-beta
HLA-B: histocompatibility antigen, B alpha chain
SRC: Proto-oncogene tyrosine-protein kinase Src
TRAF1: TNF receptor-associated factor 1
TRAF2: TNF receptor-associated factor 2
CTNNB1: Catenin beta-1
HSPA5: Endoplasmic reticulum chaperone BiP
CDKN1A: Cyclin-dependent kinase inhibitor 1
SMAD4: Mothers against decapentaplegic homolog 4
SMAD2: Mothers against decapentaplegic homolog 2
HSPB1: Heat shock protein beta-1
RELA: Transcription factor p65
PIK3R3: Phosphatidylinositol 3-kinase regulatory subunit gamma
GSK3B: Glycogen synthase kinase-3 beta
VHL: von Hippel-Lindau disease tumor suppressor
EGFR: Epidermal growth factor receptor
TP53: Cellular tumor antigen p53
PIN1: Peptidyl-prolyl cis-trans isomerase NIMA-interacting 1
SMAD3: Mothers against decapentaplegic homolog 3
KDM1A: Lysine-specific histone demethylase 1A
EEF1A1: Elongation factor 1-alpha 1
UBQLN4: Ubiquilin-4
UBC: Polyubiquitin-C
PCNA: Proliferating cell nuclear antigen
CRK: Adapter molecule crk
MDM2: E3 ubiquitin-protein ligase Mdm2
TP53: Cellular tumor antigen p53
VIM: Vimentin
MAP3K3: Mitogen-activated protein kinase 3
TNFRSF1A: Tumor necrosis factor receptor superfamily member 1A
YWHAZ: 14-3-3 protein zeta/delta
IKBKG: NF-kappa-B essential modulator
FYN: Tyrosine-protein kinase Fyn
ABL1: Tyrosine-protein kinase ABL1
GRB2: Growth factor receptor-bound protein 2
SMARCD1: SWI/SNF-related matrix-associated actin-dependent regulator of chromatin subfamily D member 1
UBE2I: SUMO-conjugating enzyme UBC9
GOLGA2: Golgin subfamily A member 2
IKBKE: Inhibitor of nuclear factor kappa-B kinase subunit epsilon
EP300: Histone acetyltransferase p300
ERBB2: Receptor tyrosine-protein kinase erbB-2
SDCBP: Syntenin-1
CDC37: Hsp90 co-chaperone Cdc37
HSCs: hepatic stellate cells

